# Longitudinal Alterations in Sleep EEG Biomarkers of Memory Consolidation in Middle-Aged and Older Adults

**DOI:** 10.64898/2026.05.11.724419

**Authors:** Destiny E. Berisha, Abhishek Dave, Negin Sattari, Miranda G. Chappel-Farley, Kate E. Sprecher, Jason Bock, Brady A. Riedner, Ethan Moran Grover, Erin M. Jonaitis, Henrik Zetterberg, Barbara B. Bendlin, Bryce A. Mander, Ruth M. Benca

## Abstract

The precise coordination of slow oscillations (SO) and sleep spindles during non-rapid eye movement (NREM) sleep supports memory consolidation and may serve as a sensitive marker of cognitive aging. However, longitudinal changes in their oscillatory dynamics in midlife and older age remain poorly understood. Using polysomnography with high-density EEG at two timepoints over ∼2.5 years, we examined changes in local NREM slow wave (SW), sleep spindle (occurring in the 11–16 Hz sigma range), and SO-sigma coupling strength in cognitively unimpaired middle-aged to older adults at risk for Alzheimer’s disease. Fronto-central SO-sigma power coupling strength significantly declined over time, independent of changes in multiple measures of SW and sleep spindle expression. Local declines in multiple sleep spindle measures were also observed. Greater baseline levels of cerebrospinal fluid (CSF) neurogranin, a postsynaptic protein abundantly expressed in the dendritic spines of the hippocampus and cerebral cortex and implicated in calcium-dependent synaptic plasticity, predicted the magnitude of longitudinal decline in SO-fast sigma coupling strength, which in turn predicted episodic memory performance changes. These findings suggest that longitudinal changes in local sleep oscillatory dynamics are related to decreased synaptic integrity and may serve as an early indicator of memory decline in older adults at risk for Alzheimer’s disease.

## Introduction

Aging is associated with alterations in global sleep architecture and local deficits in the expression of brain oscillations that define non-rapid eye movement (NREM) sleep and support neuroplasticity, e.g., slow waves and sleep spindles^1^. Cross-sectional evidence indicates these age-related oscillatory deficits are expressed locally on the scalp, with maximal observed age differences including frontal slow wave amplitude and density^2^, sleep spindle incidence, amplitude, and duration^3^, and slow wave-sleep spindle coupling strength^4^, precision^5^, and incidence^4^. However, longitudinal evidence has remained limited. A recent study reported longitudinal, frequency-specific declines in NREM spectral power over ∼5 years^6^, with decreases in 1–4 Hz slow wave activity (SWA) and 12–15 Hz sigma power exhibiting the most prominent longitudinal decline observed in middle-aged and older adults, consistent with cross-sectional findings^7–10^. Whether these longitudinal deficits are 1) local or global, 2) observed in slow wave or sleep spindle oscillatory features, or 3) associated with changes in slow wave-sleep spindle coupling, remains unclear. Moreover, the age-related mechanisms that predict these longitudinal changes in NREM sleep expression remain unknown, as do any potential functional consequences.

Cross-sectional studies provide strong evidence that aging may be associated with progressive disruption in the expression and precise coupling of NREM sleep rhythms^5,11^. Older adults also show markedly lower global absolute slow wave power, lower frontal sigma power, and lower frontal fast sleep spindle frequencies with increasing age^12^. Sleep spindle density, duration, and amplitude are also lower in older adults relative to younger adults^13,14^. Faster frequency sleep spindles (i.e., fast sigma) are also less likely to phase-lock to low frequency slow waves (termed slow oscillations, SOs), appearing to peak earlier and more variably in the SO crest, with the mean preferred phase occurring earlier relative to younger adults^5,11^. It remains unclear, however, if this coupling strength declines longitudinally, and if any such deficits are due to or are independent of changes in SO or sleep spindle expression, per se. Moreover, it is unclear what age-related factors predict longitudinal declines in sleep oscillation expression and coupling, and whether these declines predict memory trajectories.

Cross-sectional studies have shown that older adults with less structural atrophy in the medial prefrontal cortex (mPFC), thalamus, and medial temporal lobe showed more preserved NREM SWA and sleep spindle expression, and more precise, ‘youth-like’ SO-fast spindle coupling, all of which were associated with better overnight memory retention^5,10,15,16^. At the circuit level, SO-spindle (SO-SP) nesting creates a neurophysiological state characterized by maximal pyramidal cell firing activity combined with inhibitory dynamics thought to optimize synaptic plasticity within local cortical networks^17^. Given the relationship between synaptic plasticity and SOs, sleep spindles, and SO-SP coupling expression^17^, it is possible that age-related synaptic changes may predict age-related longitudinal deficits in SO-SP coupling. Furthermore, while the functional significance of these age-related sleep deficits for sleep-dependent memory processing has been reported in numerous cross-sectional studies^5,11^, it remains unclear if longitudinal declines in the expression of memory-relevant sleep oscillations contribute to trajectories of memory decline in later life. This is of particular interest, given recent findings that changes in SO-SP coupling are also linked to advancing cognitive decline and may serve as early biomarkers of Alzheimer’s disease (AD) pathology^18^. In a cohort spanning cognitively normal (CN) older adults, mild cognitive impairment (MCI) due to AD, and AD dementia, Wei and colleagues showed that SO-SP coupling becomes increasingly disrupted across the AD spectrum: coupling frequency decreased from CN to MCI, and phase locking became further misaligned in the transition from MCI to AD dementia, with disruptions associated with greater AD pathology and mPFC gray matter atrophy^18^. Importantly, SO-SP coupling misalignment predicted longitudinal decline in global cognition and episodic memory beyond other established risk factors (age, sex, education, and apolipoprotein E (*APOE*) ε4 status), and improved prediction of cognitive decline when combined with cerebrospinal fluid (CSF) and MRI biomarkers^18^. Notably, SO-SP coupling was impaired while uncoupled SOs and spindles were preserved. This suggests that impaired coupling specifically reflects degradation of memory-supporting neural coordination rather than loss of individual oscillations^18^. However, because sleep EEG was assessed at a single time point, it remains unclear whether these coupling deficits reflect stable between-person differences or progressive within-person change, and whether this degradation in coupling timing occurs independent of, or in tandem with, oscillatory loss. Consistent with this notion, older adults with early AD exhibit reduced temporal lobe sleep spindle power and density, despite preserved sleep stage proportions, which predicted faster cognitive decline over seven years. This further supports the view that disruptions in memory-relevant sleep microarchitecture reflect progressive degradation of neural systems supporting memory, which are particularly vulnerable in AD^19^.

Age-related changes in NREM SWA and sleep spindle expression are well-established features of normative aging and are associated with episodic and procedural memory impairment^1,5,11,20–22^. Mechanistically, these NREM sleep oscillations have been linked to synaptic plasticity^23^ and homeostasis^24^, as well as hippocampal–neocortical communication supporting offline sleep-dependent memory processing^25–28^. These processes are thought to be mediated by coordinated activity among cortical SOs, corticothalamic sleep spindles, and hippocampal sharp-wave ripples^27^. This coordination drives rapid increases in neuronal firing, promotes local coactivation, and facilitates spike-timing dependent synaptic plasticity and long-term potentiation^27^. At a broader level, this tripartite oscillatory coupling supports bi-directional communication between the hippocampus and neocortex, providing a supra-ordinate temporal scaffold that primes cortical circuits for the long-term storage of labile memory representations. Importantly, as spindles and SOs show marked vulnerability to aging^20^, disruption of their coordination may represent a key mechanism through which age-related sleep changes impact memory-related neural processes. As such, age-related changes in NREM sleep oscillation expression and coordination may represent a mechanistic pathway through which sleep contributes to cognitive vulnerability in aging. Understanding how these oscillations change with age, and whether they deteriorate over time within older individuals, is therefore critical for linking sleep physiology to trajectories of cognitive decline.

Despite the growing body of cross-sectional evidence on slow wave, sleep spindle, and SO-SP coupling differences with age^5,11,20,21^, several critical questions remain unresolved. Studies often utilize sparse, *a-priori* selected electrodes, limiting information on topographical changes, and no prior study has examined longitudinal changes in cross-frequency coupling of SOs and sleep spindles in middle-aged to older adults. It is unknown whether these disruptions reflect stable age-related differences or progressive within-person declines over time. It is not clear whether longitudinal changes in SO-SP coupling reflect changes in coupling strength itself or can be attributed to longitudinal changes in the expression of SOs or sleep spindles. Further, it is unknown whether changes in synaptic integrity predict vulnerability to SO-SP coupling deterioration over time. Addressing these gaps is necessary, as cross-sectional associations cannot determine whether age differences truly reflect longitudinal deterioration in sleep oscillatory dynamics or if these dynamics contribute to trajectories of cognitive decline in aging. The present study addresses these knowledge gaps by combining cerebrospinal fluid (CSF) measures of synaptic integrity, longitudinal neuropsychological testing, and longitudinal polysomnography (PSG) with high-density EEG (hdEEG) to track changes in spectral features of NREM sleep and SO-SP coupling strength in a cohort of middle-aged and older adults enriched for AD risk. We hypothesized that SO-SP coupling strength, indexed by measures of SO-sigma coupling strength, would decline longitudinally in middle-aged and older adults. We further hypothesized that baseline CSF markers of synaptic integrity would predict the magnitude of SO-fast sigma power coupling decline over time. Lastly, we explored whether local declines in SO-fast sigma were associated with differences in episodic memory trajectories.

## Methods Narrative

In short (see ***Methods*** for details), cognitively unimpaired middle-aged and older adults enriched for AD risk completed a multimodal longitudinal study combining CSF biomarkers, repeated neuropsychological testing, and overnight sleep visits. CSF was collected via lumbar puncture, after which participants completed their baseline sleep visit an average of ∼2.8 years later. They also completed a follow-up sleep visit ∼2.1 years after baseline (∼4.9 years after CSF collection). Each sleep visit included overnight PSG with concurrent high-density EEG to quantify sleep architecture, topographical spectral power, sleep spindles characteristics, and SO-sigma phase-amplitude coupling during NREM sleep. Episodic memory was assessed using the Wechsler Memory Scale–Revised (WMS-R) Logical Memory subtests across 4–11 cognitive visits spanning ∼6–12 years (mean ≈9.8 years), allowing estimation of individual trajectories of both immediate and delayed recall. Individual cognitive trajectories were estimated using linear mixed-effects models that accounted for age, sex, education, and *APOE* ε4 status, from which subject-specific slopes of annual change in immediate and delayed recall were derived. Longitudinal sleep EEG change was calculated as the annualized difference between baseline and follow-up sleep visits and evaluated with cluster-based permutation testing and mixed-effects models accounting for demographic and genetic risk factors. Associations were then examined between changes in sleep physiology, CSF biomarkers of neuronal integrity, and longitudinal trajectories of episodic memory performance.

## Online Methods

### Participants and Study Design

Participants included clinically normal, cognitively unimpaired middle-aged and older adults recruited from the Wisconsin Alzheimer’s Disease Research Center (ADRC) clinical core into a sub-study, the Predicting Alzheimer’s from Metabolic Markers and Sleep (PAMMS) study. The cohort was enriched for AD risk factors, with ∼80% reporting a parental history of AD and ∼45% carrying the *APOE* ε4 allele. Participants were recruited from the community through advertisements and word of mouth, and all were free from any significant neurological, psychiatric, or medical conditions, as well as treatments that could impact cognition, compliance with study procedures, or sleep. Use of medications known to affect sleep and sleep EEG, including antipsychotics, nonselective serotonin reuptake inhibitor (SSRI) antidepressants, neuroleptics, chronic anxiolytics, sedative–hypnotics, and stimulants, was exclusionary. Each participant completed comprehensive cognitive testing, a medical history review, and overnight polysomnography at the University of Wisconsin–Madison. Cognitive testing assessed declarative and semantic memory, attention, executive function, language, and visuospatial skills using the National Alzheimer’s Coordinating Center Uniform Data Set (UDS) neuropsychological battery version 3 alongside supplemental measures, as previously described in this cohort^29^. Memory outcomes were indexed by Logical Memory tests (see *Logical Memory*). Polysomnography (PSG) was conducted as part of a separate sub-study of 29 participants who were not currently receiving treatment for sleep-disordered breathing (e.g., CPAP) and had no major neurological, psychiatric, or medical disorders. Although obstructive sleep apnea (OSA) was not an exclusion criterion, AHI was included as a covariate in relevant analyses to account for potential effects of OSA. All participants provided written informed consent, and study protocols were approved by the University of Wisconsin–Madison Institutional Review Board.

### Polysomnography

As previously described^29^, sleep was assessed using standard polysomnography (PSG), including EOG, submental and bilateral tibial EMG, ECG, respiratory inductance plethysmography, pulse oximetry, and a position sensor, recorded with a customized Alice 5 System (Philips Respironics). High-density EEG (hdEEG) with 256 channels was concurrently recorded with vertex referencing using a NetAmps 300 amplifier and NetStation software (EGI). EEG was digitized at 500 Hz. Sleep scoring was performed using si× 10–20 hdEEG channels (F3, F4, C3, C4, O1, O2) and re-referenced to the contralateral mastoid by a registered technologist following AASM criteria^30,31^ using Alice Sleepware. A board-certified sleep physician (author R.M.B.) reviewed all scoring, including clinical sleep events. Standard sleep architecture metrics were derived, including total recording time (TRT), total sleep time (TST), sleep latency (SL), sleep efficiency (SE), wake after sleep onset (WASO), stage-specific sleep percentages (N1, N2, N3, R), apnea-hypopnea index (AHI), respiratory disturbance index (RDI), and periodic limb movement index (PLMSI). Mean values across the participant cohort for each of these variables are presented in Table 1.

**Table 1:** Participant Characteristics.

| N=29 | Baseline | Follow-Up |
| --- | --- | --- |
| Age at PSG (years, M ± SD) | 59.34 ± 4.95 | 61.48 ± 4.96 |
| BMI (M ± SD) | 27.14 ± 5.14 | 27.17 ± 4.71 |
| Sex (number) | Female – 22<br>Male – 7 |  |
| Education (years, M ± SD) | 16.76 ± 2.21 |  |
| Parental AD (number) | 23 |  |
| Presence of at least 1 APOE ε4 allele (number) | 13 |  |
| Both Parental AD and APOE ε4 (number) | 11 |  |
| Time between BASELINE, FOLLOW-UP (years, M ± SD) | 2.14 ± 0.30 |  |
| Time between earliest CSF draw and BASELINE (years, M ± SD, [range]); N=26 | 2.80 ± 1.62 |  |
| Time between earliest CSF draw and FOLLOW-UP (years, M ± SD, [range]); N=26 | 4.94 ± 1.55 |  |
| Race (number) | African American or Black - 1<br>Asian - 0<br>White - 28<br>American Indian or Alaskan Native - 0<br>More than one race - 0<br>Unknown/not reported – 0 |  |
| Ethnicity (number) | Hispanic - 1<br>Non-Hispanic – 28 |  |

### EEG Preprocessing and Multi-Taper Spectral Analysis

EEG preprocessing and spectral analyses of NREM (N2, N3) sleep were conducted in MATLAB (Mathworks, Inc., R2019b) using EEGLAB toolbox (https://sccn.ucsd.edu/eeglab/index.php) as described in our previous work^29^. Data were notch-filtered at 59.5–60.5 Hz and bandpass-filtered from 0.3–55 Hz with a 5500-point FIR filter. The outermost 50 electrodes were excluded due to artifacts, leaving 206 average-referenced channels. A three-stage semi-automated artifact rejection was applied: (1) exclusion of epochs with arousals, (2) removal of segments exceeding the 99.8th percentile of broadband power, and (3) visual rejection of residual artifacts and noisy channels. Removed channels were interpolated via spherical splines, yielding cleaned NREM data for all 29 participants. Spectral power was computed using the Chronux v2.12 toolbox (http://chronux.org) for via multi-taper analysis^32,33^ with 30s windows sliding every 5s, 11 DPSS tapers, and 0.25 Hz bandwidth. The resulting time–frequency matrix (μV²/Hz) was integrated to obtain absolute power and normalized to derive relative spectral power (unitless ratio), indicating the proportion of power in each band. For each electrode, mean absolute and relative power was calculated within canonical frequency bands were defined as follows: slow wave activity (SWA, 0.5–4.5 Hz), slow oscillations (SO, 0.5–1 Hz), delta (>1–4.5 Hz), theta (>4.5–7.5 Hz), alpha (>7.5–11 Hz), total sigma (>11–16 Hz), slow sigma (≥11–13 Hz), fast sigma (>13–16 Hz), beta (>16–28 Hz), beta1 (>16–22 Hz), beta2 (>22–28 Hz), and gamma (>28–40 Hz). Analyses focused a priori on absolute fast sigma power, motivated by prior studies demonstrating age-related changes in coupling between SOs and fast spindles^5^, the stronger involvement of SO–fast spindle coupling in memory consolidation relative to SO–slow spindle coupling5, and the selective vulnerability of fast spindle activity in the context of AD^34,35^.

### Spindle Detection

Sleep spindles were detected using the validated A7 algorithm^36^ with default thresholds, and adapted to include additional metrics as described in previous work^29^. Briefly, spindles were identified from preprocessed artifact-free N2/N3 sleep EEG across 206 channels using four parameters computed in sliding 0.3 s windows: (1) log-transformed absolute sigma power (11–16 Hz); (2) Z-normalized relative sigma power (ratio of sigma band power to broadband power, excluding delta); (3) Z-normalized sigma covariance (covariance between the broadband and sigma-filtered signals); and (4) sigma correlation (Pearson correlation between the broadband and sigma-filtered signals). Spindles were detected when all four parameters simultaneously exceeded their respective thresholds, with event durations constrained to 0.3–2.5 s. The algorithm yielded spindle count, density (count/min of artifact-free N2/N3), and duration (averaged across channels). To classify spindles as fast (≥13 Hz) or slow (<13 Hz), each detected spindle was filtered (11–<16 Hz), FFT-transformed, and labeled based on its peak spectral frequency. Mean duration and density were then calculated separately for fast and slow spindles at each channel. More details of the spindle detection approach are included in the Supplemental File.

### Phase Amplitude Coupling

As previously described^37^, slow oscillation (SO)–sigma phase amplitude coupling (PAC) was computed in two steps. First, SOs were identified from clean N2/N3 sleep EEG by detecting zero-crossings in 0.5–1 Hz filtered data. Events with negative-to-positive crossings 0.5–1.5s apart were retained if they met these criteria: ≥60 µV peak-to-peak amplitude, ≥30 µV negative half-wave depth, SO duration between 300 ms–1.5 s, and full event period ≤10 s. Next, SO and fast sigma (13-16Hz) bands were filtered with 4th and 8th/9th order Butterworth filters, respectively. The Hilbert transform was applied to extract the instantaneous SO phase and fast sigma amplitude, from which the mean vector length (MVL)^38^ and a modified modulation index (MI)^39^ were calculated. MVL quantified the strength of coupling by computing the average length of complex vectors formed by combining SO phase with spindle amplitude within time windows centered on detected slow events. The modulation index assessed the deviation of spindle amplitude distribution across SO phases from the uniform distribution, using Kullback-Leibler divergence with bins spanning the full cycle (−π to π). For both metrics, surrogate distributions were generated via circular shifting of the amplitude signal to derive normalized (z-scored) values and permutation-based p-values; normalized values were used in all analyses to account for subject- and channel-specific variability in amplitude and phase structure. The preferred phase of coupling (i.e., the SO phase at which spindle amplitude peaked) was estimated from the angle of the complex MVL vector. All analyses were restricted to local epochs surrounding detected SOs to isolate physiologically meaningful coupling and reduce confounding from non-oscillatory segments. This algorithm was repeated for SO-slow sigma (11-13Hz) coupling as well for completeness, with findings reported in Supplementary Materials. SO density, number, and amplitude were also extracted from this algorithm.

### Logical Memory

Logical Memory A-I (LM A-I) and Logical Memory A-II (LM A-II) are subtests from the Weschler Memory Scale Revised (WMS-R; as previously described^40^), designed to assess episodic memory using two short narratives (“A” and “B”), each comprising 25 informational “idea units.” Each story is read aloud to the participant, who is then asked to recall it immediately after presentation (LM A-I, immediate recall) and again after a 25–30 minute delay without re-exposure (LM A-II, delayed recall). Participants may recall the items in any order. Scoring follows WMS-R manual guidelines, which permit certain paraphrases (e.g., “slid off the table” for “fell off the table”) while requiring verbatim recall for some content such as proper names and numerical expressions. For each participant, immediate and delayed recall scores are calculated as the total number of idea units recalled across both stories, averaged over Stories A and B. Both LM A-I and LM A-II are sensitive to early episodic memory decline, which represents the earliest and most severely affected cognitive domain in AD and a critical marker for distinguishing amnestic mild cognitive impairment from normal aging^41^.

### CSF Collection and Analysis

Cerebrospinal fluid (CSF) collection, processing, and biomarker quantification were performed as previously described^29^. In brief, CSF was collected in the morning after an overnight fast via lumbar puncture at the L3/4 or L4/5 interspace using a 24- or 25-gauge Sprotte needle. Approximately 22 mL of CSF was gently extracted, mixed to prevent gradient effects, centrifuged at 2000 × g for 10 minutes, aliquoted (0.5 mL), and stored at −80 °C within 30 minutes. Assays were performed by blinded, board-certified technicians. Biomarker quantification was performed with Roche NeuroToolKit, an investigational panel of automated immunoassays, run on cobas-e-601 and e-411 analyzers at the Clinical Neurochemistry Laboratory, University of Gothenburg. Analytes included the axonal injury marker neurofilament light chain (NfL) and synaptic proteins neurogranin and α-synuclein.

### Statistical Analysis

#### Longitudinal Changes in Sleep EEG Measures

To compare changes from baseline to follow-up, an annualized change matrix was calculated. Briefly, a matrix was aggregated containing data from each subject at each electrode using the median value across the time series data, resulting in a subject by electrode matrix (subjects x electrodes, for baseline and follow-up, separately). These matrices were then subtracted, and the resulting matrix was divided row-wise by an array of years-between-visits corresponding to time between baseline and follow-up for each subject ((FOLLOWUP - BASELINE)/Years). This resultant matrix was then used in a paired t-test against a dimension-matched matrix of zeros followed by 5000-permutation threshold free cluster enhancement (TFCE) to reveal significant changes at the electrode level. To confirm that these changes were robust against covariates (sex, *APOE* ε4 carrier status, age at baseline visit, and log-transformed AHI), a mixed effects linear model was implemented to assign random effects as the subject-level variability in a data frame in which *time* was a column with entries ‘visit 1’ and ‘visit 2’ to contain data from both visits. The models were implemented such that the target measure in the significant electrodes resulting from the TFCE-multiple comparison correction was assigned as the dependent variable as a multivariate model with each significant electrode detected from TFCE. Parental AD and education years were not included as covariates as they were non-significant in each model (p>0.1) and thus removed to improve the power of the models. For coupling measures relating to phase angle, circular statistics were used to compare phase angle matrices at baseline and follow-up, where Watson-Wheeler tests were applied to evaluate differences in angular distributions (MATLAB, CircStat P. Berens, CircStat: A MATLAB Toolbox for Circular Statistics, Journal of Statistical Software, Volume 31, Issue 10, 2009 http://www.jstatsoft.org/v31/i10). For multiple comparison corrections not involving spectral matrix data, p-values were corrected for multiple comparisons using the Benjamini–Hochberg False Discovery Rate (FDR) procedure where appropriate^42^. In correlational analyses, Pearson correlations were computed to test associations for variables that met the assumptions of normality; otherwise, a Kendall’s τ correlation was used.

#### Cluster Analyses

Cluster analyses were accomplished by averaging the annualized changes of spectral measures in the largest-detected cluster of electrodes that significantly changed over time in the whole sample (see Figure S1). The largest clusters in the TFCE-corrected annualized change in EEG measures were detected using a graph-based approach based on spherical distances of electrodes. In short, electrode locations were converted to unit vectors, and pairwise angular distances were computed to form adjacency matrices connecting electrodes within a proximity threshold. Connected components were identified using MATLAB’s *graph* and *conncomp* functions, and the largest component was retained as the primary spatial cluster, with both clusters being in fronto-central regions of the EEG topography for each coupling metric (Figure S1).

#### CSF Analyses

Levels of CSF biomarkers were examined prior to the baseline sleep visit (Table S1): participants completed their first sleep visit an average of 2.80 years after CSF collection (range: 0.43–6.33 years; SD = 1.62) and their second sleep visit an average of 4.94 years after CSF collection (range: 2.32–8.61 years; SD = 1.55). Subjects with CSF data yielded a subset of N = 26. Three CSF biomarkers were included in this analysis that reflect synaptic/neuronal integrity: neurogranin (log-transformed), NfL, and α-synuclein (reciprocal).

#### Logical Memory Slopes

To estimate the rate of change in Logical Memory performance (immediate: LM A-I and delayed: LM A-II) for each participant, separate linear mixed-effects models were fit for each outcome using the lme4 package in R (Bates et al., 2015). Each model included fixed effects for years since first cognitive test, age at earliest cognitive test, sex, *APOE* ε4 carrier status, and years of education, as well as random intercepts and random slopes for years since first testing by participant. Only participants with at least three visits (N = 28) were included. The range of visits used for this cohort were between four and 11 visits, with a mean of 7.76±1.66 visits. Years from first test to last follow-up were an average of 9.83 ±1.65 years, with a range of six to 12 years. For each participant, the subject-specific slope for years since first testing was extracted from the model’s conditional coefficients. These slopes represent the estimated annual change in raw score units for each recall type, adjusted for timing, demographic and genetic risk covariates. Negative slopes indicate a decline in recall performance per year, whereas positive slopes indicate improvement.

## Results

### Participant Characteristics

Twenty-nine cognitively unimpaired middle-aged and older adults completed two PSG studies with hdEEG (baseline PSG visit mean age±SD: 59.342 ± 4.948 years; age at follow-up PSG: 61.5 ± 5.0 years; 22 female, baseline AHI 6.8 ± 10.2; Table 1). Relative to the general population, this cohort was enriched for AD risk factors, with 13 *APOE* ε4 carriers and 23 participants with a parental history of AD. The mean time between PSG visits was 2.1±0.3 years. Details of sleep architecture for each PSG visit are reported in Table 2, with no significant differences in sleep architecture variables detected between visits (all p>0.05). A subset of 26 participants consented to CSF collection, which occurred on average 2.8±1.6 years before the baseline PSG visit. One participant was amyloid positive, as indicated by a Aβ 42/40 ratio threshold of 0.041, and none exhibited tau positivity based on validated thresholds^43^.

**Table 2:** Sleep Reports.

| Measure | Baseline | Follow-Up |
| --- | --- | --- |
| *Sleep Architecture* |  |  |
| **Total Sleep Time (TST, min)** | 373.67 ± 63.07 | 369.93 ± 88.51 |
| **Time in Bed (TIB, min)** | 448.24 ± 60.61 | 462.50 ± 77.28 |
| **Wake After Sleep Onset (WASO, min)** | 60.69 ± 39.64 | 72.32 ± 39.91 |
| **Sleep Efficiency (SE, %)** | 83.49 ± 10.24 | 78.86 ± 17.15 |
| **Sleep Latency (SL, min)** | 13.88 ± 16.65 | 20.25 ± 28.99 |
| **REM Time (min)** | 75.91 ± 28.52 | 72.71 ± 28.79 |
| **REM Percentage (R, %)** | 19.92 ± 5.41 | 18.68 ± 6.09 |
| **REM Latency (REML, min)** | 105.19 ± 54.24 | 115.13 ± 56.67 |
| **NREM Time (min)** | 297.76 ± 46.39 | 297.39 ± 69.45 |
| **N1 Percentage (N1, %)** | 6.55 ± 5.87 | 7.73 ± 9.12 |
| **N1 Time (N1, min)** | 24.31 ± 22.40 | 22.70 ± 14.40 |
| **N2 Percentage (N2, %)** | 58.92 ± 10.44 | 56.85 ± 8.16 |
| **N2 Time (N2, min)** | 220.21 ± 53.53 | 212.43 ± 62.32 |
| **N3 Percentage (N3, %)** | 14.61 ± 9.94 | 16.73 ± 8.30 |
| **N3 Time (N3, min)** | 53.57 ± 38.54 | 62.43 ± 28.85 |
| **Wake Time (W, min)** | 74.24 ± 47.52 | 92.41 ± 50.32 |
| *Sleep Disorder Variables* |  |  |
| **AHI** | 6.8 ± 10.21 | 5.49 ± 10.17 |
| **Arousals Index (events/hr)** | 18.00 ± 11.33 | 21.10 ± 15.75 |
| **Time <90 SpO2 (mins)** | 2.60 ± 5.43 | 7.73 ± 21.63 |
| **Mean SpO2 (%)** | 95.21 ± 1.32 | 95.00 ± 1.33 |
| **Min SpO2 (%)** | 87.17 ± 5.575 | 87.93 ± 5.07 |
| **PLMSI TST (events/hr)** | 15.66 ± 17.24 | 12.16 ± 14.11 |

### Longitudinal Change in Local Sleep in Midlife and Old Age

#### Changes in Slow Oscillation and Sleep Spindle Expression

We first sought to characterize local longitudinal changes in NREM SOs and SPs. To assess this, topographical annualized changes in relative power within SO (0.5-1 Hz) and fast sigma (13-16 Hz) frequency bands were assessed. Significant changes were detected in relative fast sigma across frontal, occipital, and temporal regions following TFCE correction, Figure 1A. Following the detection of significant electrodes with TFCE in relative fast sigma power only, a subsequent model adjusting for covariates revealed a significant longitudinal decline in relative fast sigma power (β=-0.00003, p<0.001), with age at baseline additionally predicting lower relative fast sigma power cross-sectionally (β=-0.000005, p=0.026; Suppl. Table 1C). No significant changes were detected in relative SO power, Figure S2E (middle panel). Analysis of changes in absolute power are shown in Figure S3. Analysis of longitudinal change in absolute and relative power in other frequency bands are shown in Figure S4, with notable global reductions in relative beta power (16-28 Hz frequency band) and sparse reductions in absolute theta power (4.5-7.5 Hz).

**Figure 1.**
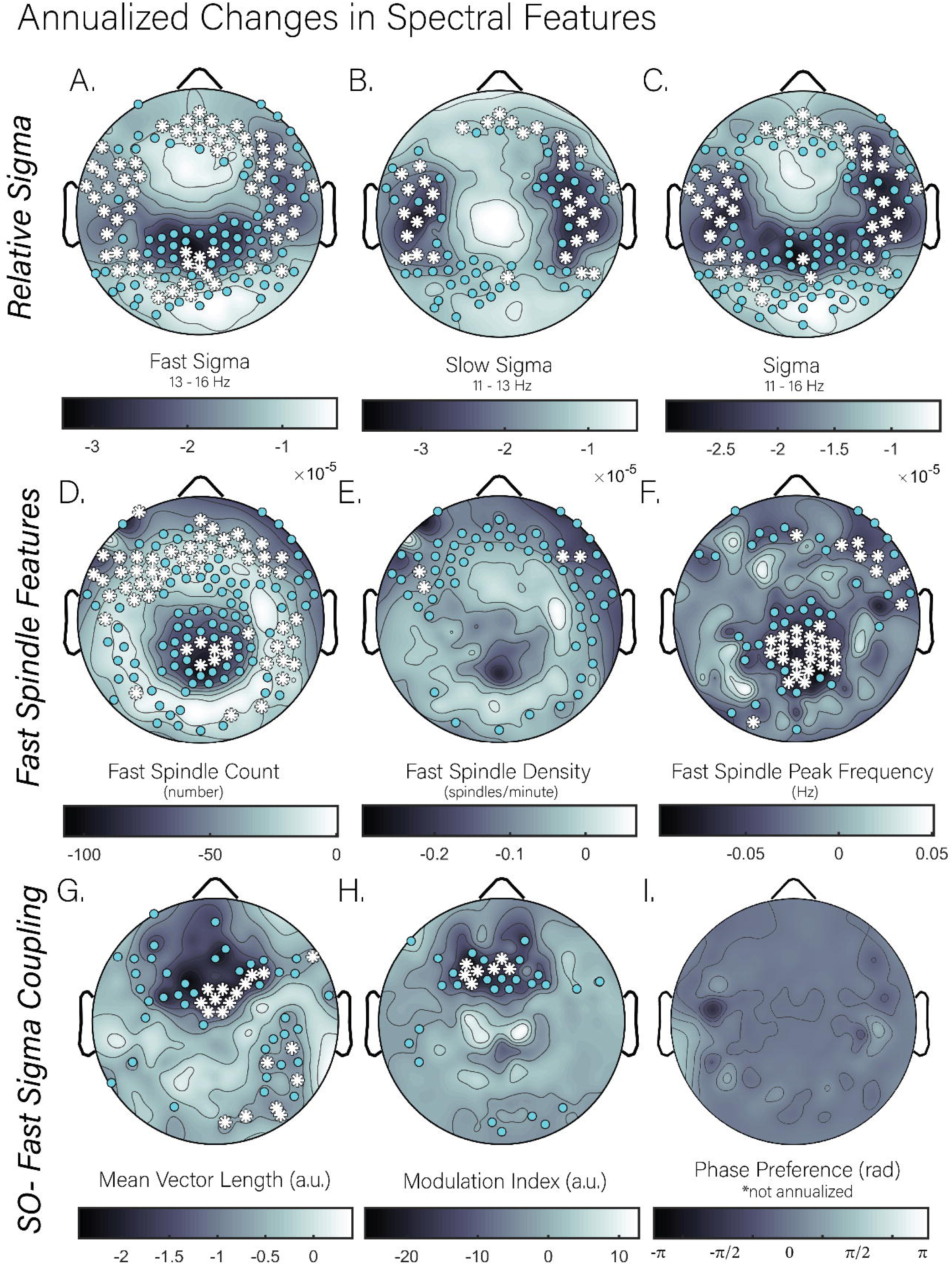
Topographic maps of annualized change in spectral features from baseline to follow-up. The top row reflects annualized changes in relative fast sigma (1A; 13-16 Hz), slow sigma (1B; 11-13 Hz), and total sigma (1C; 11-16Hz) power. The middle row shows annualized changes in fast spindle features including count (1D), density (1E), and peak frequency (1F). Bottom row: The first two panels show annualized changes in fast spindle-slow oscillation coupling metrics: mean vector length and modulation index, respectively (1G-H). Panel 3 (right; 1C) shows the circular difference in mean coupling phase between visits (follow-up subtracted by baseline, not annualized), with values wrapped to the range (–π, π); positive values indicate a later phase preference, and negative values indicate an earlier preference.

For completeness, models characterizing longitudinal changes in relative delta (1-4.5 Hz) and total sigma (11-16 Hz), and slow sigma (11-13 Hz) were also examined (respectively, Figures S2F, 1C, and 1B). Models adjusting for covariates (Suppl. Table 1D) revealed a significant longitudinal decline in relative delta power (β = -0.00015, p < 0.001). In addition, there were independent, cross-sectional effects detected, such that older age at baseline (β = 0.000022, p = 0.045) and higher AHI (β = 0.000064, p = 0.038) both predicted lower relative delta. A similar model for relative total sigma power similarly revealed a significant longitudinal decline in relative total sigma power (β=-0.00003, p<0.001, Suppl. Table 1E) and in relative slow sigma power (β=-0.00003, p<0.001, Suppl. Table 1I).

As relative power in sigma frequency bands does not necessarily reflect sleep spindle changes alone, fast sleep spindle features (i.e., peak frequency, count, density, and duration) were examined using an established sleep spindle algorithm validated in older adults.^36^ This was used to determine whether similar longitudinal declines were observed in sleep spindle characteristics. Analysis of annualized change revealed significant and broad declines over frontal and central derivations following TFCE correction in all measures but fast sleep spindle duration, Figure 1D-F. In mixed models accounting for time and covariates, all sleep spindle features with significant electrodes following TFCE correction (density, peak frequency, and count) showed a robust and significant decrease with time (density: β=-0.23, p<0.001; peak frequency: β=-0.05, p<0.001; count: β=-73.0, p<0.001; Tables 3F-H respectively). Higher log-AHI was also significantly associated with higher peak frequency cross-sectionally (β=0.03, *p*<0.001). SO characteristics were also evaluated (count, density, amplitude), but no features exhibited significant longitudinal change over 2 years (all p>0.05, Figure S2A-C).

#### Changes in Slow Oscillation- Fast Sigma Coupling

We next sought to characterize longitudinal changes in the coordination between SOs and fast sigma power. One sample t-tests of annualized change in mean vector length (MVL) and modulation index (MI) revealed longitudinal decreases in SO-fast sigma coupling at fronto-central electrodes following TFCE correction, with reduced MVL also detected at right temporal derivations, Figure 1G and 1H. However, as for mean SO phase of fast sigma coupling, no longitudinal change was detected, Figure 1I. Side-to-side comparisons of SO-fast sigma coupling and SO-slow sigma coupling are depicted in Figure S5 for completeness, demonstrating a more robust decline in SO-fast sigma coupling than SO-slow sigma coupling, consistent with previous cross-sectional findings expected^5^.

Significant clusters showing TFCE significance in longitudinal changes in MVL and MI were extracted and included in subsequent mixed-effects models adjusting for age, sex, *APOE* ε4 status, and AHI covariates to determine if the observed longitudinal decreases were robust to inclusion of covariates. These models revealed a significant longitudinal reduction in both MVL (β=–1.24, *p*<0.001, Suppl. Table 1A) and MI (β= –19.70, *p*<0.001, Suppl. Table 1B) measures at follow-up compared to baseline. Independent, cross-sectional covariate effects on SO-fast sigma coupling were also observed, with higher log-AHI significantly predicting higher raw MVL (β=1.11, *p*<0.001); however, no evidence indicated that AHI predicted longitudinal change in coupling measures (all p>0.05) when MVL clusters were assessed with baseline AHI and change in AHI (follow-up – baseline; Figure S 6). No effects of age or *APOE* ε4 carrier status were observed for MVL or MI in these models. These findings indicate that SO-fast sigma coupling decreases longitudinally in middle aged and older adults, particularly over fronto-central EEG derivations.

Given that fast sleep spindle expression declined over the study period, it is possible that the decline in SO-fast sigma coupling is secondary to disruptions in sleep spindle expression. To explore this possibility, we tested whether longitudinal changes in slow wave and sleep spindle characteristics were associated with longitudinal changes in SO-fast sigma coupling measures. We extracted largest cluster of significant TFCE-corrected electrodes showing longitudinal decreases in MVL and MI for these analyses (Figure S1). Longitudinal change in the MVL cluster showed no significant correlations with longitudinal change in fast sleep spindle peak frequency (r=0.16, p=0.23), fast sleep spindle count (r=0.07, p=0.60), absolute or relative sigma power (|r|<0.11, all p>0.44), or SO power (absolute: r=-0.19, p=0.32; relative: r=-0.23, p=0.22). Similarly, change in MI was generally not significantly associated with any change in spectral power or sleep spindle features, although a trend-level correlation emerged with fast spindle count (r=0.25, p=0.066). These findings suggest that changes in sleep spindles and SOs over time do not robustly explain the longitudinal decrease in SO-fast sigma coupling in middle-aged and older adults.

### Baseline Markers of Synaptic Integrity & Memory Decline

Since prior studies have demonstrated that SO-SP coupling is associated with synaptic plasticity^17,27^ and structural integrity^11,20^, we sought to determine whether severity of synaptic degeneration might predict longitudinal changes in SO-fast sigma coupling. Annualized change in MI was significantly and negatively correlated with log-transformed CSF Neurogranin levels (Kendall’s τ=–0.342, FDR-corrected p = 0.042; Figure 2), indicating that lower synaptic integrity at baseline predicted greater reductions in SO-fast sigma coupling strength over time. No significant associations were observed between MI and CSF NfL (Kendall’s τ=0.009, uncorrected p=0.965, FDR-corrected p=0.965) or α-Synuclein (reciprocal-transformed; Kendall’s τ=0.120, uncorrected p=0.406, FDR-corrected p=0.609). To determine whether the relationship between CSF neurogranin levels and change in MI remained significant after correcting for covariates, a multiple regression model was implemented adjusting for age at baseline PSG visit, sex, and the time difference between the follow-up PSG visit and the CSF draw, which revealed that CSF neurogranin levels (log-transformed) significantly predicted MI (β=-63.63, *p*=0.034). No significant associations were observed with other baseline CSF markers and MVL (all FDR-corrected p>0.255). These findings suggest that a lower synaptic integrity may be associated with greater progressive decoupling of memory-relevant sleep oscillations over time.

**Figure 2:**
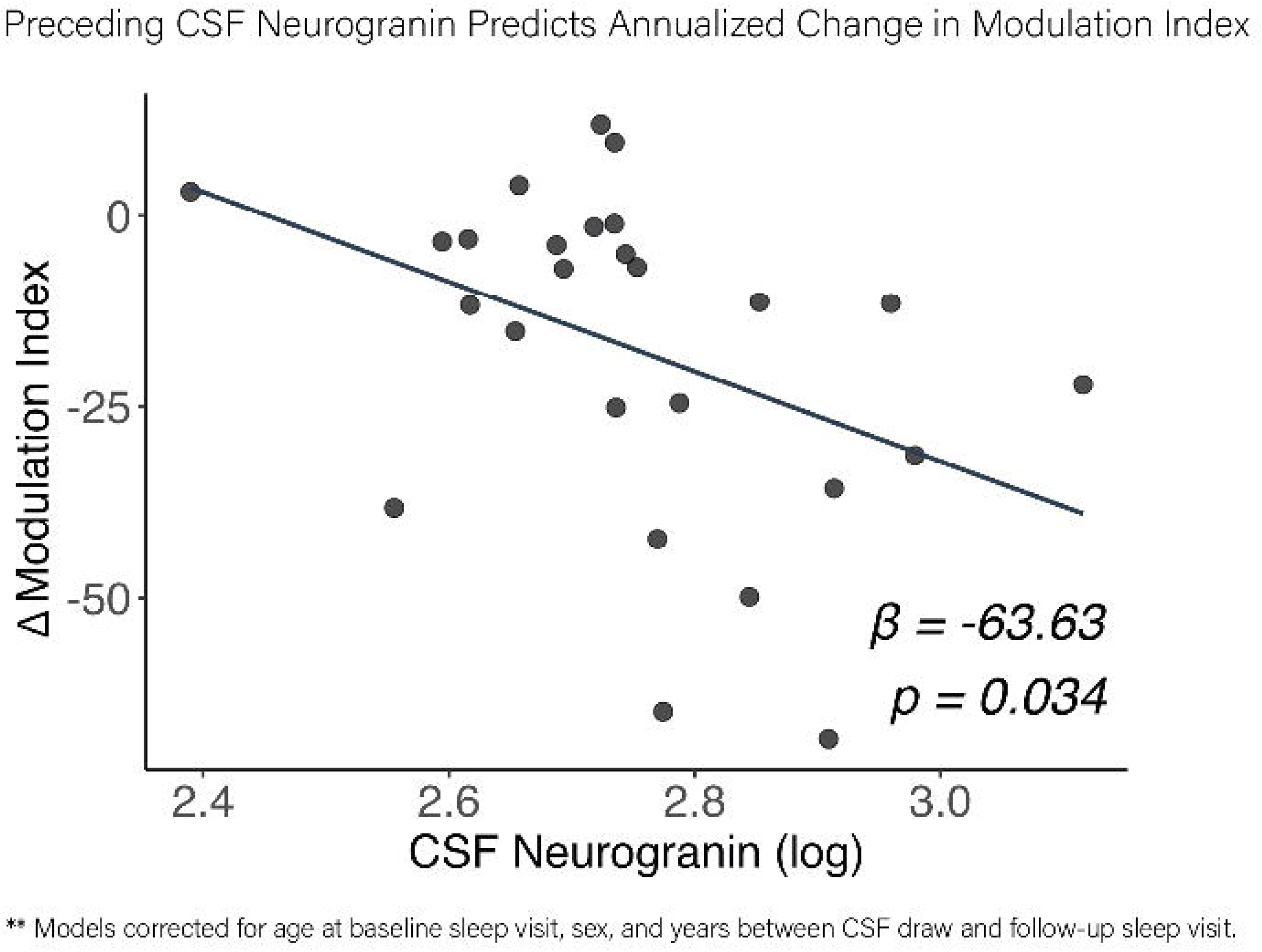
Preceding CSF Neurogranin Predicts Annualized Change in Modulation Index of SO-Fast Sigma. Association of CSF neurogranin (log-transformed for normality) at baseline with annualized change in modulation index at the largest detected cluster of TFCE-corrected electrodes, adjusting for age at baseline PSG visit, sex, and time difference in years between CSF draw and the follow-up sleep visit.

To explore whether the annualized decline in coupling predicted changes in episodic memory performance, associations were examined between slopes of delayed-recall Logical Memory scores across at least three visits per participant. Annualized change in MVL was associated with delayed recall slopes (r = 0.39, p=0.038, Figure 3). No association was observed for MI (Kendall’s τ = 0.06, p>0.1). For an additional control analysis, immediate recall slopes were not associated with either MVL or MI (respectively r=-0.069, Kendall’s τ=-0.025, both p>0.1). Baseline levels of CSF neurogranin were not significantly associated with delayed recall slopes (r=0.198, p>0.05).

**Figure 3:**
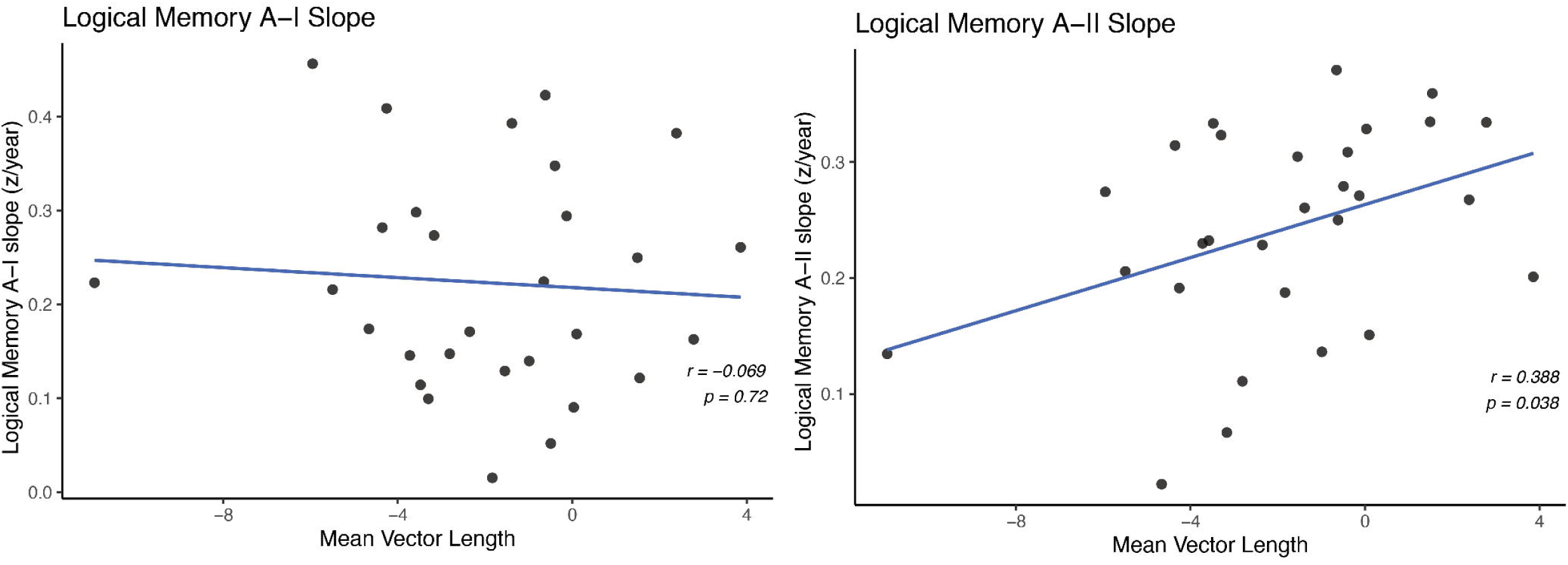
Annualized change in Mean Vector Length of SO-Fast Sigma Coupling Predicts Delayed-Recall Memory Slopes. Left: association of annualized change in mean vector length to rate of change in immediate recall of logical memory. Right: association of annualized change in mean vector length to rate of change in delayed recall of logical memory. Rate of change in logical memory performance (immediate: LM A-I and delayed: LM A-II) for each participant was determined by fitting linear mixed-effects models with years since first testing, age at earliest test, sex, *APOE* ε4 carrier status, and years of education, as well as random intercepts and random slopes for years-since-first-testing by participant. Participants with at least three visits (N = 28) were included. For each participant, the subject-specific slope for years since first testing from the model’s conditional coefficients was extracted and used as the slope. Negative slopes indicate a decline in recall performance per year, whereas positive slopes indicate improvement.

## Discussion

Consistent with prior cross-sectional findings^11,13^, SO-fast sigma coupling strength and sleep spindle expression measures exhibited regional longitudinal declines over ∼2.5 years. These deficits were larger in individuals with higher CSF-measured levels of synaptic dysfunction at baseline. Moreover, exploratory analyses indicated that individuals with greater longitudinal reductions in frontal SO-fast sigma coupling exhibited faster deterioration in episodic memory. Collectively, these findings suggest sleep oscillatory mechanisms supporting memory processing weaken over time and that the synaptic dysfunction may contribute to this process. Specifically, reduced SO-fast spindle coupling over frontal cortex may reflect disruption of hippocampal–neocortical communication essential to memory function, contributing to episodic memory decline.

Longitudinal changes in sleep oscillatory dynamics, particularly the coupling of SOs and fast spindles in middle- to older-aged adults remain understudied, despite growing recognition of its relevance for cognitive aging and neurodegenerative processes^18,29,44,45^, and this age range being considered an optimal prodromal window for intervention. Most existing studies have relied on cross-sectional data and pre-selected electrodes, limiting insight into individual trajectories and scalp-wide patterns^5,6,11^. This study addresses these limitations by leveraging longitudinal PSG with hdEEG recordings to examine within-subject changes in NREM spectral features and SO-fast sigma coupling in cognitively at-risk middle-aged and older adults. In this cohort, there was a robust longitudinal decline in SO-fast sigma coupling, which was not explained by concurrent declines slow wave or sleep spindle features and power. This supports the hypothesis that the de-coupling of fast spindles from SOs may be mechanistically distinct from disruptions in the expression of individual sleep oscillations in older adults.

The consistent cross-sectional association between frontal gray matter structural integrity and SO-SP coupling from prior studies^11,18^ motivated our examination of CSF markers of synaptic integrity in relation to longitudinal changes in coupling, as the frontal cortex shows some of the earliest and most pronounced age-related structural decline, much of which is thought to reflect synaptic loss and remodeling^46^. Previously reported associations between frontal gray matter volume and SO-SP coupling may reflect underlying synaptic integrity, leading to our focus on CSF biomarkers of synaptic and neuronal health. We found that greater declines in SO-fast sigma coupling strength were predicted by higher baseline levels of CSF neurogranin, a protein found in postsynaptic dendritic spines^47^. In contrast, CSF α-synuclein, a pre-synaptic protein regulating neurotransmitter release^48^, did not independently predict coupling decline in this cohort, nor did CSF neurofilament light chain protein (NfL), a marker of axonal injury^49^. Taken together, these findings suggest that age-related changes in SO-fast sigma coupling may be more closely tied to postsynaptic dendritic alterations characteristic of normative aging, rather than to overt neurodegeneration or neuronal loss. Given that SO-SP coupling reflects precisely timed interactions among distributed neural populations^27^, it is likely to be particularly sensitive to age-related reductions in synaptic integrity. Notably, neurogranin binds calmodulin and is thought to modulate synaptic plasticity and long-term potentiation (LTP) through the regulation of calcium/calmodulin-dependent signaling pathways^50,51^ and is abundantly expressed in the brain, particularly within the dendritic spines of the hippocampus and cerebral cortex: regions involved in the generation of oscillations critical for memory consolidation during sleep. Therefore, increased synaptic dysfunction in these regions may gradually impair pathways critical for information transfer within the mPFC–thalamic–hippocampal circuit, yielding progressive SO-SP decoupling that deteriorates memory processing over time. In this context, CSF neurogranin may represent a more sensitive marker of aging-related synaptic vulnerability than other CSF markers of axonal damage or neurodegeneration. Given the specific role of SO-fast spindle coupling in memory consolidation, we suspected that changes in coupling would predict cognitive trajectories in this cohort. Indeed, our exploratory findings support this possibility, with annualized change in MVL predicting slopes of episodic memory performance in this cohort.

Surprisingly, unlike prior cross-sectional and longitudinal studies in middle aged and older adults^6–9^, we did not find robust declines in SWA expression in this cohort, apart from a few electrodes showing significant reductions in relative delta power. No significant changes were detected in the SO features and SO relative power, consistent with Gao and colleagues’ findings in a large longitudinal cohort of middle-aged and older adults^6^. The reported decline in sigma power across central channels and in relative delta in a few channels is also consistent^6^. Given collective these trends, we can surmise that fast spindle activity may be especially vulnerable to normative aging while SOs may be relatively spared in the absence of an active neurodegenerative process. Notably, cross-sectional work comparing young and middle-aged adults shows that aging is associated more so with altered slow-wave dynamics (lower frequency, longer positive and negative phase durations, and reduced slope, especially over prefrontal/frontal regions) than uniform reductions in slow-wave power^2^, which aligns with the relative stability of SO power observed here. However, the lack of robust SWA power reductions in our cohort may also reflect the relatively short follow-up interval (∼2.5 years), suggesting that measurable SWA decreases may emerge over a longer time frame.

Our *a priori* focus on fast spindles was motivated by past studies examining age-related changes in coupling of SOs to fast spindles^5^, stronger evidence implicating fast spindle coupling to SO’s is most important for memory consolidation processes relative to coupling with slow spindles^5^, and their specific vulnerability to aging^13,14^ and AD^34,35^. That features of fast spindles changed over time was not surprising. With the decline in fast spindle peak frequency over time, it is possible that across the lifespan, fast spindles may shift to slower frequencies. This might explain why in one study, SO-slow SP coupling proportion was similar in young adults and older adults^5^. As slow spindles tend to be coupled to the up-to-down state transition of SOs, a period characterized by gradual neuronal silence, SO-slow SP coupling may be less relevant for memory consolidation processes, a hypothesis supported by evidence in younger adults^5^ but not in older adults^52^. Similarly, a study of healthy older adults demonstrated that a slow spindle coupling phase closer to the SO up-state in a frontal EEG channel was predictive of better memory^52^, supporting the notion that the precise phase of coupling matters more than the type of spindle in older adults. In contrast to prior cross-sectional findings^11,21^, we did not detect a longitudinal change in the SO phase angle at which fast sigma power tended to peak. It is possible that the duration of the follow-up period and the small sample size were insufficient to detect this change. It is also possible this age effect is not apparent in longitudinal data. Future studies in larger samples with longer follow-up periods will be important to disentangle these possibilities.

While this study has many strengths (high density EEG, whole-scalp analysis, and a highly characterized cohort), it is not without limitations. First, the average time between visits was only ∼2.5 years, which precluded longer-term assessments of longitudinal change in the NREM spectral features. Second, data about use and compliance of CPAP was not available for these participants, which may have confounded analyses of sleep-disordered breathing on longitudinal changes in the coupling metrics and other spectral features. Third, this sample was relatively small, largely White, and female, limiting generalizability to more diverse populations. This is of note, given that a prior longitudinal study of spectral power during sleep measured at a central EEG channel found sex-specific cross-sectional age associations, with males showing lower NREM power across multiple frequency bands, including SO, delta, and sigma, whereas females primarily exhibited age-related reductions in sigma power, despite no longitudinal change in NREM SO power^6^. Given these findings, it is possible that coupling decreases in men might be further exacerbated by decreases in SWA. Future studies with larger, sex-balanced cohorts should explore longitudinal changes in coupling to elucidate this further.

Together, our findings support a unified account in which age-related decoupling of SO-fast sigma coordination reflects a network-level vulnerability that is partly independent of global changes in slow wave and sleep spindle expression. Specifically, we show that (i) fast spindle expression and SO-fast sigma coupling strength decline longitudinally, (ii) a marker of synaptic integrity, CSF neurogranin, predicts the magnitude of SO-fast sigma decoupling over time, and (iii) longitudinal decline in SO-fast sigma coupling is associated with the longitudinal trajectory of episodic memory decline. We propose a mechanistic framework in which aging and decreased synaptic integrity together degrade mPFC–thalamic–hippocampal crosstalk, yielding progressive SO-SP decoupling that in turn weakens memory consolidation abilities over time. The functional consequence of decreased SO-fast sigma coupling may be mitigated by neurostimulation to enhance SO-SP coupling^53^. For example, in a recent intracranial EEG study of adults, stimulation of the prefrontal cortex during SO up-states in the medial temporal lobe enhanced SO-SP–ripple coupling across regions and improved declarative recognition memory^53^. While brain stimulation holds promise as a potential intervention to support memory in aging, its efficacy and reproducibility are yet to be fully established. Beyond aging, stimulation to enhance SO-SP coupling may also reduce cognitive symptom burden in individuals across the AD disease spectrum.

## Abbreviations

AD: Alzheimer’s disease
SO: Slow Oscillation
SP: Sleep Spindle
SO-SP: Slow Oscillation–Sleep Spindle (Coupling)
MVL: Mean Vector Length
MI: Modulation Index
TFCE: Threshold-Free Cluster Enhancement
AHI: Apnea-Hypopnea Index
REM: Rapid Eye Movement
OSA: Obstructive Sleep Apnea
PSG: Polysomnography
CSF: Cerebrospinal Fluid
NfL: Neurofilament Light Chain
α-Synuclein: Alpha-Synuclein
*APOE* ε4: Apolipoprotein E epsilon 4 allele
FDR: False Discovery Rate
mPFC: Medial Prefrontal Cortex
SWS: Slow Wave Sleep
SWA: Slow Wave Activity

## Article Information

### Author Contributions

D.E.B. analyzed the data and wrote the manuscript. Authors A.D., N.S.S., and M.G.C-F aided in writing programming/code algorithms to generate the EEG variables, author K.E.S collected the data, author J.B. consulted on statistical approaches, author B.A.R. aided in pre-processing the data, author E.M.G. aided in data management, author E.M.J. aided in memory outcomes harmonization, author H.Z. aided in CSF measures, authors B.B.B., B.A.M., and R.M.B. edited the manuscript and study concept or design.

### Conflict of Interest Disclosures

Dr. Mander has served as a consultant for Eisai Co., Ltd and currently serves on a scientific advisory board for AstronauTx. Dr. Benca has served as a consultant to Eisai, Genentech, Haleon, Idorsia, and Sage, and has received research grant support from Eisai. Dr. Chappel-Farley has served as a consultant to Apnimed. Dr. Zetterberg has served at scientific advisory boards and/or as a consultant for Abbvie, Acumen, Alamar, Alector, Alzinova, ALZpath, Amylyx, Annexon, Apellis, Artery Therapeutics, AZTherapies, Cognito Therapeutics, CogRx, Denali, Eisai, Enigma, LabCorp, Merck Sharp & Dohme, Merry Life, Nervgen, Novo Nordisk, Optoceutics, Passage Bio, Pinteon Therapeutics, Prothena, Quanterix, Red Abbey Labs, reMYND, Roche, Samumed, ScandiBio Therapeutics AB, Siemens Healthineers, Triplet Therapeutics, and Wave, has given lectures sponsored by Alzecure, BioArctic, Biogen, Cellectricon, Fujirebio, LabCorp, Lilly, Novo Nordisk, Oy Medix Biochemica AB, Roche, and WebMD, is a co-founder of Brain Biomarker Solutions in Gothenburg AB (BBS), which is a part of the GU Ventures Incubator Program, and is a shareholder of CERimmune Therapeutics (outside submitted work).

### Funding Information

This project was made possible by the following grants: R56 AG052698, R01 AG027161, R01 AG021155, ADRC P50 AG033514, R01 AG037639, K01 AG068353, and National Research Service Awards F31 AG084308 and F31 AG048732 from the National Institute on Aging, and by the Clinical and Translational Science Award (CTSA) program, through the NIH National Center for Advancing Translational Sciences (NCATS), grant UL1TR000427. Dr. Zetterberg is a Wallenberg Scholar supported by grants from the Swedish Research Council (#2018-02532), the European Research Council (#681712), Swedish State Support for Clinical Research (#ALFGBG-720931), the Alzheimer Drug Discovery Foundation (ADDF), USA (#201809-2016862), the AD Strategic Fund and the Alzheimer’s Association (#ADSF-21-831376-C, #ADSF-21-831381-C and #ADSF-21-831377-C), the Olav Thon Foundation, the Erling-Persson Family Foundation, Stiftelsen för Gamla Tjänarinnor, Hjärnfonden, Sweden (#FO2019-0228), the European Union’s Horizon 2020 research and innovation programme under the Marie Skłodowska-Curie grant agreement No 860197 (MIRIADE), and the UK Dementia Research Institute at UCL. Author HZ is a Wallenberg Scholar and a Distinguished Professor at the Swedish Research Council supported by grants from the Swedish Research Council (#2023-00356, #2022-01018 and #2019-02397), the European Union’s Horizon Europe research and innovation programme under grant agreement No 101053962, and Swedish State Support for Clinical Research (#ALFGBG-71320).

### Additional Contributions

We would like to thank participants and staff of the Wisconsin ADRC and Wisconsin Sleep for their contributions to the study, and the laboratory technicians at the Clinical Neurochemistry Laboratory, in Mölndal, Sweden. Without their efforts this research would not be possible. CSF assays kits were provided by Roche Diagnostics GmbH. The content is solely the responsibility of the authors and does not necessarily represent the official views of the NIH.

### Data Sharing Statement

See Supplement.

