## Supplemental Files for "Longitudinal Alterations in Sleep EEG Biomarkers of Memory Consolidation in Middle-Aged and Older Adults"

**Study Type:** Research Article, Observational Study

Authors:

Destiny E. Berisha^1,2^, Abhishek Dave^3,4^, Negin Sattari^4^, Miranda G. Chappel-Farley^6^, Kate E. Sprecher^7-9^, Jason Bock^1,2^

Brady A. Riedner^10^, Ethan Moran Grover^11^, Erin M. Jonaitis^12^, Henrik Zetterberg^13-17^, Barbara B. Bendlin^9,18,19^, Bryce A. Mander^2,4,5,20,21^, Ruth M. Benca^22^

1 Department of Neurobiology and Behavior, University of California, Irvine, CA, USA

2 Center for the Neurobiology of Learning and Memory, University of California, Irvine, CA, USA

3 Department of Cognitive Sciences, University of California, Irvine, CA, USA

4 Department of Psychiatry and Human Behavior, University of California, Irvine, CA, USA

5 Institute for Memory Impairments and Neurological Disorders, University of California, Irvine, CA, USA

6 Department of Psychiatry, University of Pittsburgh School of Medicine, Pittsburgh, PA, USA

7 Neuroscience Training Program, University of Wisconsin–Madison, Madison, WI, USA

8 Department of Medicine, University of Wisconsin–Madison, Madison, WI, USA

9 Wisconsin Alzheimer’s Disease Research Center, University of Wisconsin–Madison, Madison, WI, USA

10 Department of Psychiatry, University of Wisconsin–Madison, Madison, WI, USA

11 University of Wisconsin–Madison, Madison, WI, USA

12 School of Nursing, University of Wisconsin–Madison, Madison, WI, USA

13 Institute of Neuroscience and Physiology, University of Gothenburg, Gothenburg, Sweden

14 Clinical Neurochemistry Laboratory, Sahlgrenska University Hospital, Mölndal, Sweden

15 Department of Neurodegenerative Disease, UCL Institute of Neurology, Queen Square, London, UK

16 UK Dementia Research Institute at UCL, London, UK

17 Department of Pathology and Laboratory Medicine, University of Wisconsin School of Medicine and Public Health, Madison, WI, USA

18 Wisconsin Alzheimer’s Institute, Madison, WI, USA

19 Geriatric Research Education and Clinical Center, Wm. S. Middleton Veterans Hospital, Madison, WI, USA

20 Department of Pathology and Laboratory Medicine, University of California, Irvine, CA, USA

21 Department of Neurology, University of California, Irvine, CA, USA

22 Department of Psychiatry and Behavioral Medicine, Wake Forest University, Winston-Salem, NC, USA

****Corresponding Authors**

Bryce A. Mander, Ph.D.

5141 California Ave, Room 238, Irvine CA, 92617

949-824-6742

Ruth M. Benca, M.D., Ph.D.

791 Jonestown Road

Winston Salem, NC 27103

(336) 716-2911;

Destiny E. Berisha, Ph.D.

1416 Biological Sciences III, Irvine CA, 92697

646-269-5549

**Supplementary Table 1.** Linear Mixed Effects Models Assessing Changes Over Time in Spectral Measures

| **A) *Mean Vector Length*** | β | *SE* | *t* | *p* |
| --- | --- | --- | --- | --- |
| Intercept | 12.178 | 8.493 | 1.434 | 0.164 |
| Time (Visit 2) | -1.241 | 0.259 | -4.802 | <.001 *** |
| Sex (Male) | 1.108 | 1.657 | 0.669 | 0.510 |
| APOE ε4 (Carrier) | 0.036 | 1.414 | 0.026 | 0.980 |
| Age at Baseline | -0.179 | 0.143 | -1.251 | 0.223 |
| AHI (log-transformed) | 1.109 | 0.247 | 4.483 | <.001 *** |
| *Random Effects:* | *Group* | *Variance* | *SD* |  |
| No. Obs: 1023 | *Subjects* | *13.500* | *3.674* |  |
| Groups: 29, Subjects | *Residual* | *15.690* | *3.961* |  |

| **B) *Modulation Index*** | β | *SE* | *t* | *p* |
| --- | --- | --- | --- | --- |
| Intercept | 132.704 | 110.088 | 1.205 | 0.239 |
| Time (Visit 2) | -19.700 | 2.239 | -8.797 | <.001 *** |
| Sex (Male) | -7.704 | 21.335 | -0.361 | 0.721 |
| APOE ε4 (Carrier) | 6.913 | 18.311 | 0.378 | 0.709 |
| Age at Baseline | -1.214 | 1.854 | -0.655 | 0.519 |
| AHI (log-transformed) | -0.728 | 2.195 | -0.332 | 0.740 |
| *Random Effects:* | *Group* | *Variance* | *SD* |  |
| No. Obs: 324 | *Subjects* | *2308.900* | *48.050* |  |
| Groups: 29, Subjects | *Residual* | *433.800* | *20.830* |  |

| **C. *Relative Fast Sigma Power*** | β | *SE* | *t* | *p* |
| --- | --- | --- | --- | --- |
| Intercept | 4.550e-04 | 1.172e-04 | 3.882 | <.001 *** |
| Time (Visit 2) | -2.992e-05 | 2.055e-06 | -14.555 | <.001 *** |
| Sex (Male) | 2.054e-05 | 2.268e-05 | 0.906 | 0.374 |
| APOE ε4 (Carrier) | -3.568e-05 | 1.949e-05 | -1.831 | 0.079 . |
| Age at Baseline | -4.687e-06 | 1.974e-06 | -2.375 | 0.026 * |
| AHI (log-transformed) | 1.618e-06 | 2.025e-06 | 0.799 | 0.424 |
| *Random Effects:* | *Group* | *Variance* | *SD* |  |
| No. Obs: 3564 | *Subjects* | *2.627e-09* | *5.125e-05* |  |
| Groups: 29, Subjects | *Residual* | *3.444e-09* | *5.868e-05* |  |

| **D. *Relative Delta Power*** | β | *SE* | *t* | *p* |
| --- | --- | --- | --- | --- |
| Intercept | 2.116e-03 | 6.186e-04 | 3.421 | <0.01 ** |
| Time (Visit 2) | -1.529e-04 | 3.478e-05 | -4.396 | <.001 *** |
| Sex (Male) | 1.293e-04 | 1.218e-04 | 1.062 | 0.298 |
| APOE ε4 (Carrier) | 7.674e-05 | 1.020e-04 | 0.752 | 0.459 |
| Age at Baseline | 2.181e-05 | 1.029e-05 | 2.119 | 0.045 * |
| AHI (log-transformed) | 6.363e-05 | 3.034e-05 | 2.097 | 0.039 * |
| *Random Effects:* | *Group* | *Variance* | *SD* |  |
| No. Obs: 108 | *Subjects* | *6.388e-08* | *0.0003* |  |
| Groups: 29, Subjects | *Residual* | *3.034e-08* | *0.0002* |  |

| **E. *Relative Sigma Power*** | β | *SE* | *t* | *p* |
| --- | --- | --- | --- | --- |
| Intercept | 5.300e-04 | 1.695e-04 | 3.128 | <0.01 ** |
| Time (Visit 2) | -3.457e-05 | 2.442e-06 | -14.155 | <.001 *** |
| Sex (Male) | 1.764e-05 | 3.229e-05 | 0.546 | 0.590 |
| APOE ε4 (Carrier) | -4.484e-05 | 2.778e-05 | -1.614 | 0.119 |
| Age at Baseline | -5.767e-06 | 2.814e-06 | -2.049 | 0.051 . |
| AHI (log-transformed) | -7.952e-07 | 2.416e-06 | -0.329 | 0.742 |
| *Random Effects:* | *Group* | *Variance* | *SD* |  |
| No. Obs: 2646 | *Subjects* | *5.356e-09* | *7.318e-05* |  |
| Groups: 29, Subjects | *Residual* | *3.608e-09* | *6.007e-05* |  |

| **F. *Fast Spindle Density (/min)*** | β | *SE* | *t* | *p* |
| --- | --- | --- | --- | --- |
| Intercept | 2.517 | 1.299 | 1.938 | 0.064 . |
| Time (Visit 2) | -0.233 | 0.055 | -4.202 | <.001 *** |
| Sex (Male) | -0.076 | 0.257 | -0.296 | 0.770 |
| APOE ε4 (Carrier) | -0.282 | 0.216 | -1.306 | 0.204 |
| Age at Baseline | -0.028 | 0.022 | -1.293 | 0.208 |
| AHI (log-transformed) | -0.018 | 0.052 | -0.345 | 0.731 |
| *Random Effects:* | *Group* | *Variance* | *SD* |  |
| No. Obs: 202 | *Subjects* | *0.301* | *0.549* |  |
| Groups: 29, Subjects | *Residual* | *0.142* | *0.376* |  |

| **G. *Fast Spindle Peak Freq. (Hz)*** | β | *SE* | *t* | *p* |
| --- | --- | --- | --- | --- |
| Intercept | 13.950 | 0.285 | 48.941 | <.001 *** |
| Time (Visit 2) | -0.049 | 0.005 | -10.269 | <.001 *** |
| Sex (Male) | 0.063 | 0.055 | 1.149 | 0.261 |
| APOE ε4 (Carrier) | -0.060 | 0.047 | -1.270 | 0.216 |
| Age at Baseline | -0.003 | 0.005 | -0.639 | 0.529 |
| AHI (log-transformed) | 0.029 | 0.005 | 6.042 | <.001 *** |
| *Random Effects:* | *Group* | *Variance* | *SD* |  |
| No. Obs: 2037 | *Subjects* | *0.016* | *0.125* |  |
| Groups: 29, Subjects | *Residual* | *0.011* | *0.104* |  |

| **H. *Fast Spindle Count*** | β | *SE* | *t* | *p* |
| --- | --- | --- | --- | --- |
| Intercept | 623.194 | 315.678 | 1.974 | 0.060 . |
| Time (Visit 2) | -72.999 | 8.005 | -9.120 | <.001 *** |
| Sex (Male) | -24.563 | 61.530 | -0.399 | 0.693 |
| APOE ε4 (Carrier) | -71.439 | 52.526 | -1.360 | 0.186 |
| Age at Baseline | -6.782 | 5.318 | -1.275 | 0.214 |
| AHI (log-transformed) | 8.143 | 7.759 | 1.049 | 0.294 |
| *Random Effects:* | *Group* | *Variance* | *SD* |  |
| No. Obs: 2860 | *Subjects* | *18789* | *137.1* |  |
| Groups: 29, Subjects | *Residual* | *42153* | *205.3* |  |

| **I. *Relative Slow Sigma*** | β | *SE* | *t* | *p* |
| --- | --- | --- | --- | --- |
| Intercept | 7.578e-04 | 2.436e-04 | 3.110 | 0.005 ** |
| Time (Visit 2) | -3.856e-05 | 3.595e-06 | -10.725 | <.001 *** |
| Sex (Male) | 2.488e-05 | 4.709e-05 | 0.528 | 0.602 |
| APOE ε4 (Carrier) | -4.427e-05 | 4.051e-05 | -1.093 | 0.285 |
| Age at Baseline | -7.595e-06 | 4.103e-06 | -1.851 | 0.076 . |
| AHI (log-transformed) | -4.523e-06 | 3.556e-06 | -1.272 | 0.204 |
| *Random Effects:* | *Group* | *Variance* | *SD* |  |
| No. Obs: 1890 | *Subjects* | *1.139e-08* | *1.067e-04* |  |
| Groups: 29, Subjects | *Residual* | *5.584e-09* | *7.473e-05* |  |

**Supplementary Table 1.** Linear mixed-effects models assessing longitudinal change in sleep spectral and spindle measures. Separate models were fit for each outcome (A–I). Fixed effects included time (follow-up “visit 2” vs. baseline “visit 1”), sex, APOE ε4 carrier status, age at baseline, and log-transformed apnea–hypopnea index. Random intercepts were specified for subject. β = regression coefficient; SE = standard error; t = t-statistic; p = p-value. AHI = apnea–hypopnea index; APOE = apolipoprotein E; Hz = hertz. Statistical significance: p < .05 (), p < .01 (), p < .001 (); trend-level: p < .10 (.).

**Supplementary Figure 1:**

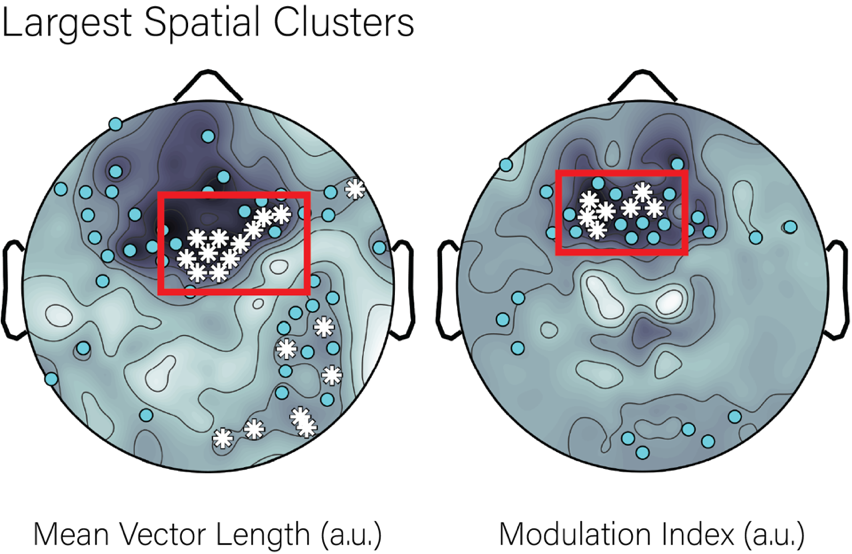

**Figure S1:** Two examples of the resultant largest clusters in the TFCE-corrected annualized change in EEG measures, highlighted in red. Clusters were detected using a graph-based approach based on spherical distances of electrodes. In short, electrode locations were converted to unit vectors, and pairwise angular distances were computed to form adjacency matrices connecting electrodes within a proximity threshold. Connected components were identified using MATLAB’s graph and *conncomp* functions, and the largest component was retained as the primary spatial cluster.

**Supplementary Figure 2:**

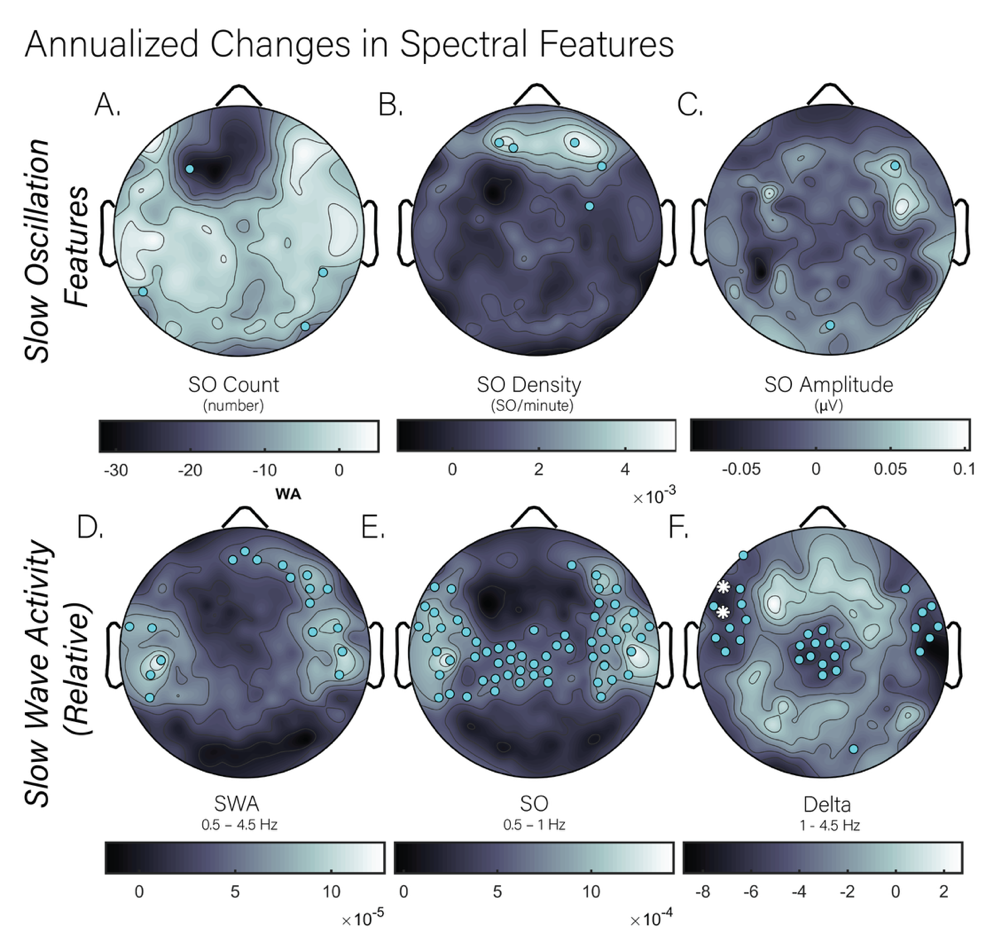

**Figure S2:** Annualized changes in slow oscillation features are shown in the top row including slow oscillation (SO) count (left), SO density (middle), and SO amplitude (right). The bottom row shows annualized change in relative SWA activity, including relative total SWA (left; 0.5-4.5 Hz), slow oscillation power (0.5-1 Hz), and delta power (1-4.5 Hz).

**Supplementary Figure 3:**

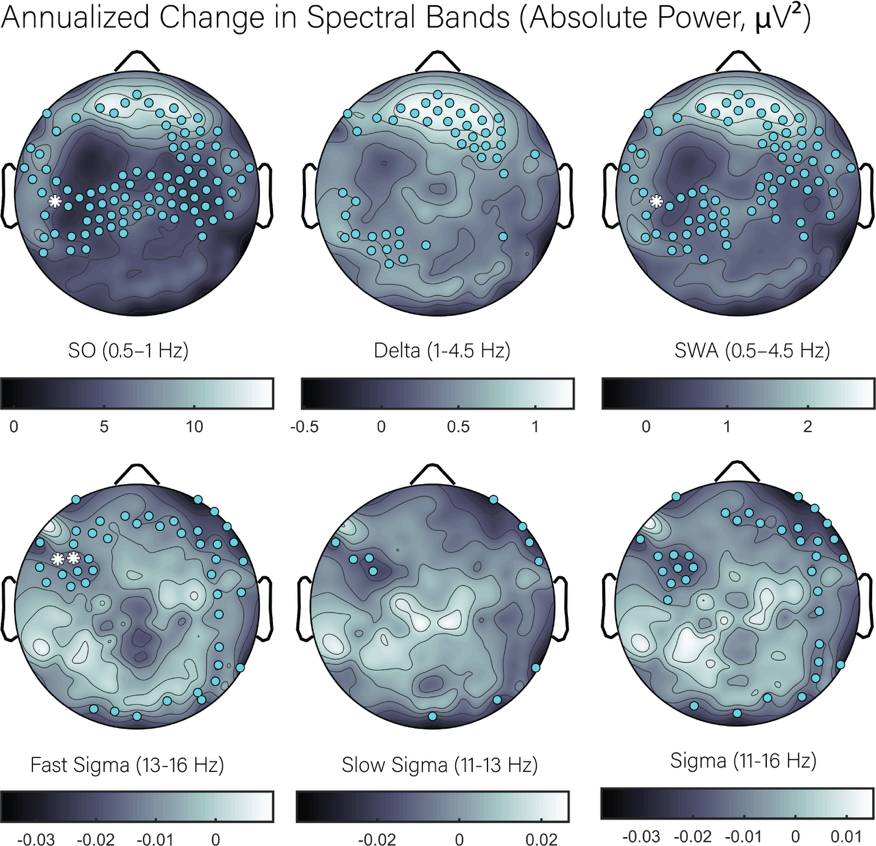

**Figure S3:** Absolute changes in frequency bands of interest. The top row displays annualized changes in absolute power of the slow wave activity (SWA) bands: slow oscillations (left; 0.5-1 Hz), delta (middle; 1-4.5 Hz), and all SWA (0.5-4.5 Hz). The bottom row shows annualized changes in absolute power across sigma bands: fast sigma (left; 13-16 Hz), slow sigma (middle; 11-13 Hz), and total sigma (right; 11-16 Hz).

**Supplementary Figure 4:**

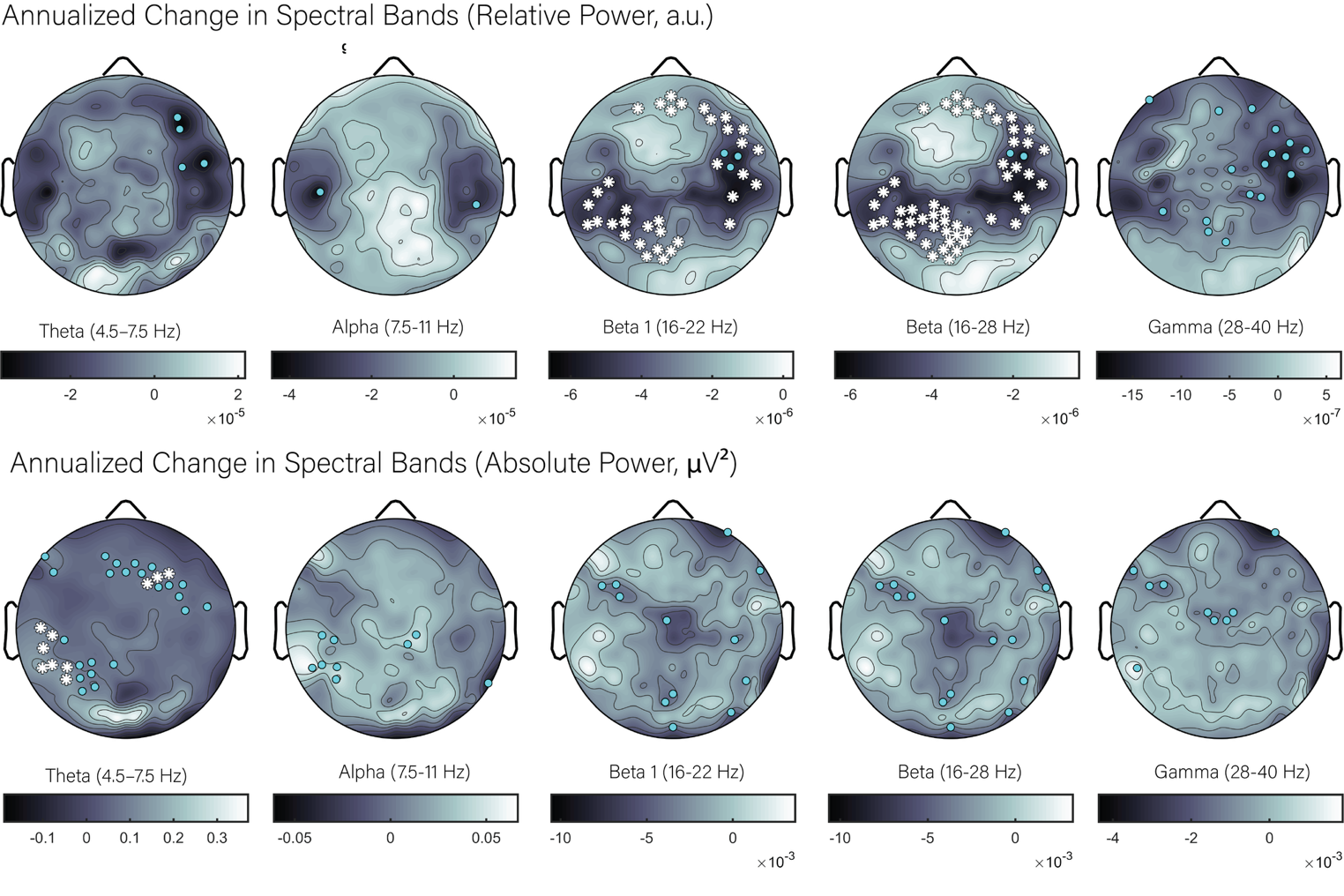

**Figure S4:** Changes in relative (top row) and absolute (bottom row) power are shown. For both rows, left to right, are shown annualized changes in theta (4.5-7.5 Hz), alpha (7.5-11 Hz), beta 1 (16-22 Hz), total beta (16-28 Hz), and gamnma (28-40 Hz) power.

**Supplementary Figure 5:**

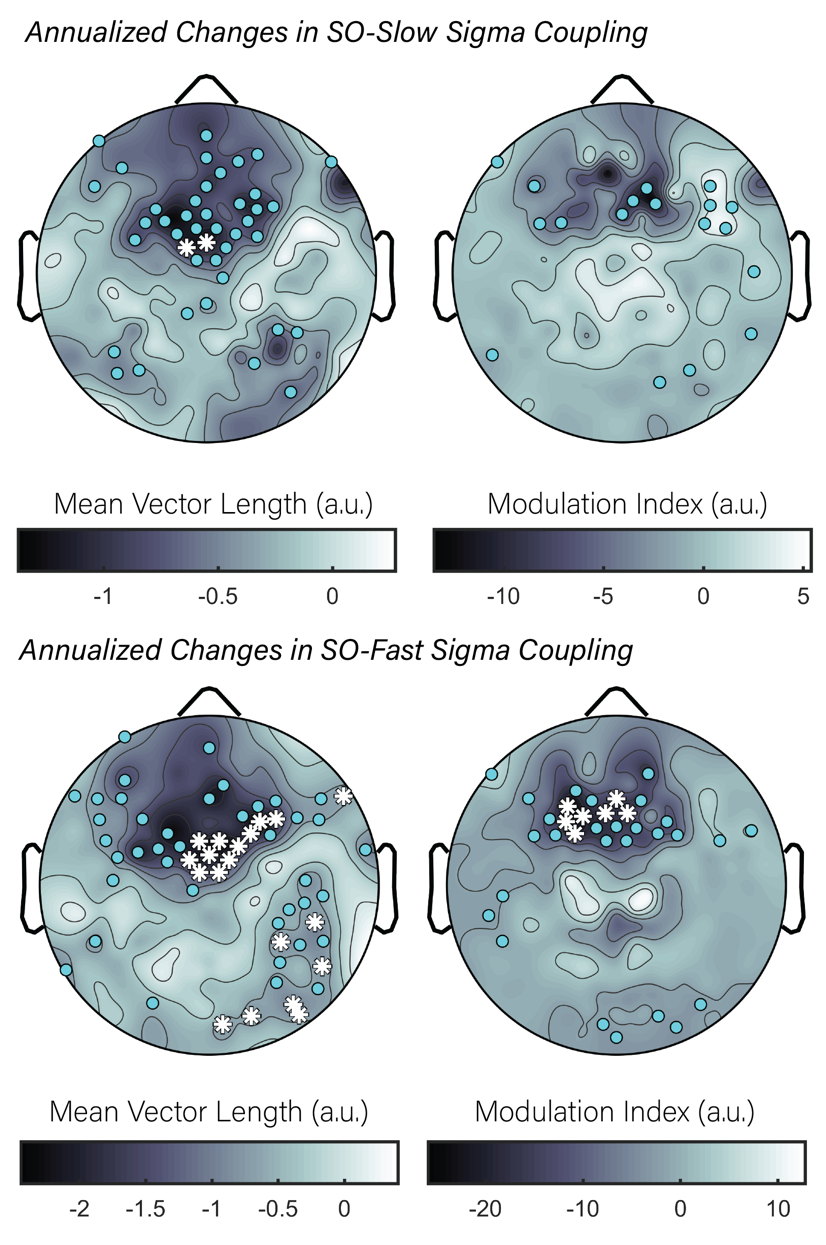

**Figure S5:** Side-by-side comparisons of annualized changes in SO-slow sigma (top row) and SO-fast sigma coupling (bottom row) are shown. Left panels show the Mean Vector Length, and the right shows Modulation Index.

**Supplementary Figure 6:**

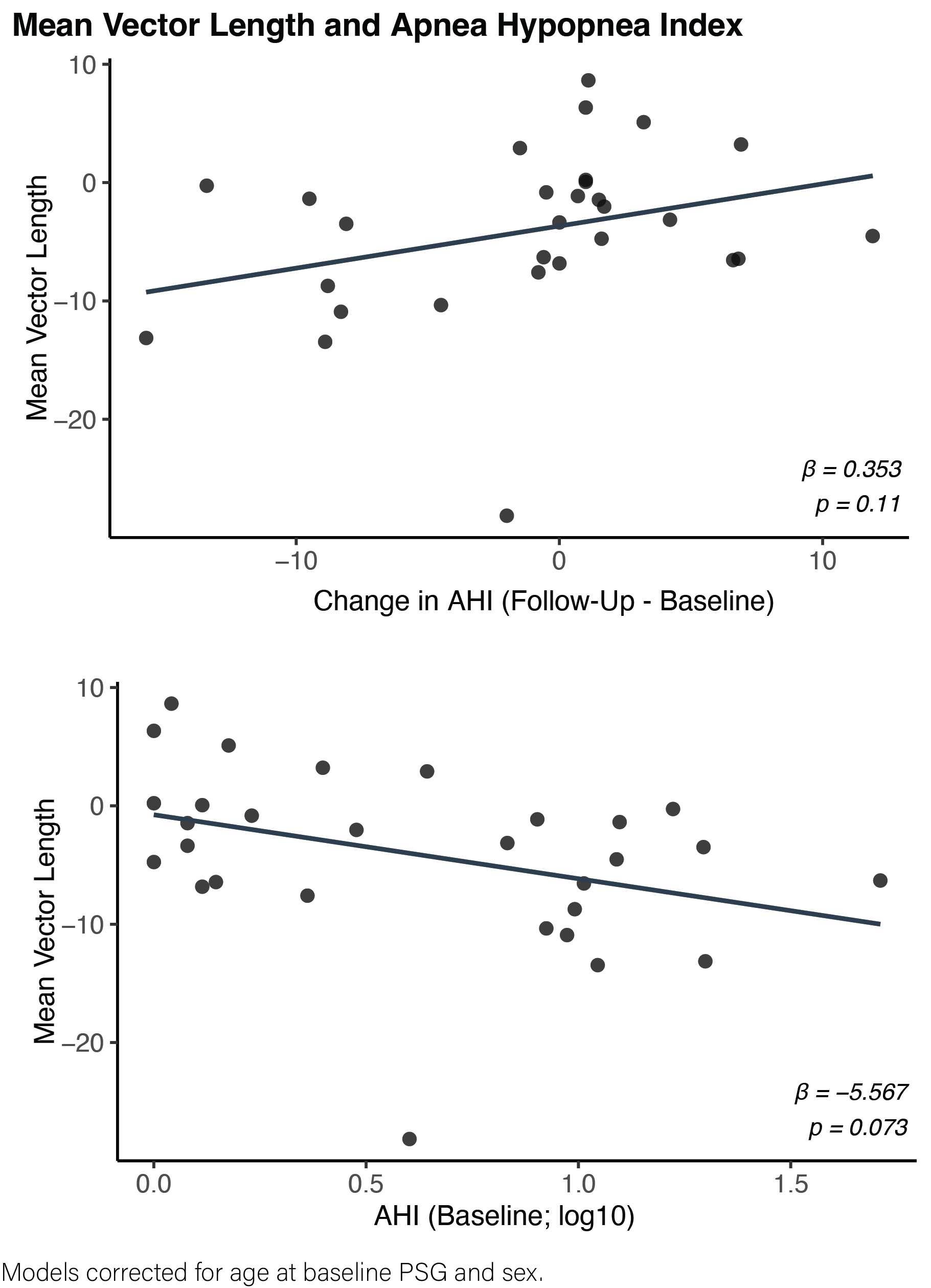

**Figure S6:** Analysis of change in Apnea-Hypopnea Index (AHI) from baseline to follow-up (top row) against changes in mean vector length (MVL) from baseline to follow-up, and analysis of baseline AHI to change in MVL from baseline to follow-up.

**Supplementary Table 2:**

| **Variable** | **CSF Draw**  Mean±SD [Min, Max] |
| --- | --- |
| CSF age (years) | 56.392±4.645 [47.870, 65.470] |
| α-Synuclein (pg/mL) | 116.419±41.715 [69.620, 254.300] |
| Aβ-40 (pg/mL) | 11808.846±3179.086 [6900.000, 18350.000] |
| Aβ-42 (pg/mL) | 785.612±261.344 [405.100, 1373.000] |
| Neurogranin (pg/mL) | 596.908±221.137 [245.300, 1306.000] |
| NfL (pg/mL) | 64.699±22.431 [31.230, 118.800] |
| Aβ42/40 ratio | 0.066±0.010 [0.041, 0.084] |
| p-Tau/Aβ42 ratio | 0.019±0.007 [0.010, 0.046] |
| t-Tau/Aβ42 ratio | 0.213±0.072 [0.120, 0.486] |

**Supplementary Table 2.** Cerebrospinal fluid (CSF) biomarker characteristics at time of lumbar puncture. Values are presented as mean ± standard deviation with range [minimum, maximum]. CSF = cerebrospinal fluid; Aβ = amyloid-β; GFAP = glial fibrillary acidic protein; IL-6 = interleukin-6; NfL = neurofilament light chain; p-Tau = phosphorylated tau; t-Tau = total tau; sTREM2 = soluble triggering receptor expressed on myeloid cells 2; YKL-40 = chitinase-3-like protein 1 (CHI3L1). Ratios represent biomarker concentration ratios as indicated.

**Supplementary Table 3:**

| **A) *Mean Vector Length*** | β | *SE* | *t* | *p* |
| --- | --- | --- | --- | --- |
| Intercept | 11.887 | 8.563 | 1.388 | 0.177 |
| Time (Visit 2) | -0.523 | 0.464 | -1.128 | 0.260 |
| Sex (Male) | 1.142 | 1.670 | 0.684 | 0.500 |
| AHI (log trans.) | 1.288 | 0.265 | 4.855 | <.001 *** |
| APOE ε4 (Carrier) | 0.019 | 1.426 | 0.013 | 0.990 |
| Age at Baseline | -0.179 | 0.144 | -1.238 | 0.227 |
| Time(Visit2) x AHI(log trans.) | -0.504 | 0.271 | -1.865 | 0.063 . |
| *Random Effects:* | *Group* | *Variance* | *SD* |  |
| No. Obs: 1023 | *Subjects* | *13.72* | *3.705* |  |
| Groups: 29, Subjects | *Residual* | *15.65* | *3.956* |  |

| **B) *Modulation Index*** | β | *SE* | *t* | *p* |
| --- | --- | --- | --- | --- |
| Intercept | 128.886 | 110.207 | 1.169 | 0.253 |
| Time (Visit 2) | -10.253 | 3.993 | -2.568 | 0.011 * |
| Sex (Male) | -7.244 | 21.354 | -0.339 | 0.737 |
| AHI (log trans.) | 1.599 | 2.323 | 0.688 | 0.492 |
| APOE ε4 (Carrier) | 6.666 | 18.329 | 0.364 | 0.719 |
| Age at Baseline | -1.208 | 1.856 | -0.651 | 0.521 |
| Time(Visit2) x AHI(log trans.) | -6.628 | 2.330 | -2.845 | < 0.01 ** |
| *Random Effects:* | *Group* | *Variance* | *SD* |  |
| No. Obs: 378 | *Subjects* | *2314.3* | *48.11* |  |
| Groups: 29, Subjects | *Residual* | *425.1* | *20.62* |  |

**Supplementary Table 3.** Linear mixed-effects models testing the interaction between time and sleep apnea severity on spectral coupling measures. Separate models were fit for each outcome (A–B). Fixed effects included time (Visit 2 vs. baseline), sex, APOE ε4 carrier status, age at baseline, log-transformed apnea–hypopnea index (AHI), and the Time × AHI interaction. Random intercepts were specified for subject. β = regression coefficient; SE = standard error; t = t-statistic; p = p-value; AHI = apnea–hypopnea index; APOE = apolipoprotein E. Statistical significance: p < .05 (), p < .01 (), p < .001 (); trend-level: p < .10 (.).

**Methods:**

*Spindle Detection*

Sleep spindles were detected and quantified using the validated A7 algorithm relying on default threshold settings, as described previously^1^. In short, sleep spindles were detected in artifact-free, concatenated signals in N2 and N3 staged epochs in the same 206 hdEEG channels used for spectral analysis. The A7 algorithm uses a combination of log-transformed absolute power, Z-normalized log-transformed relative power, and the correlation and covariance between the original N2 and N3 time series and the bandpass filtered signal in the range 11–<16 Hz. These four parameters are all computed in 0.3 s sliding time windows with a 0.1 s step size. Each parameter is associated with a detection threshold, and all four detection thresholds must be exceeded simultaneously for underlying data to be classified as belonging to a sleep spindle. Targeted spindle durations were in the range 0.3–2.5 s. Outputs generated from the detection algorithm included sleep spindle count, sleep spindle density (spindle count divided by minutes of artifact-free N2/N3 sleep), and sleep spindle durations. Sleep spindle durations were then averaged across channels to yield mean spindle duration for each of the 206 electrodes.

To additionally classify detected sleep spindles as either fast (13–<16 Hz) or slow (11–<13 Hz) spindles, previously detected spindles were extracted individually from cleaned NREM data that were filtered in the 11–<16 Hz range, for each subject at each electrode. Every sleep spindle then underwent an FFT-based spectral power calculation, using a rectangular window with zero-padding to the nearest power of 2. The peak frequency in each spindle segment was identified as the frequency corresponding to the highest value of spectral power. Sleep spindles were classified as either fast or slow based on their calculated peak frequencies, with peak frequencies below 13 Hz corresponding to slow sleep spindles and peak frequencies at or above 13 Hz corresponding to fast sleep spindles. Subsequently, the mean fast and slow sleep spindle durations were calculated separately for each subject at each EEG derivation by averaging the durations of individual fast and slow sleep spindles. Fast and slow sleep spindle densities were calculated by dividing the count of fast and slow sleep spindles respectively by minutes of artifact-free N2 and N3 sleep stages.
